# ACKR1-expressing venous endothelial cells establish a pro-fibrotic niche in pulmonary fibrosis

**DOI:** 10.64898/2026.07.31.742106

**Authors:** Konstantinos Kontodimas, Ahmed A. Raslan, Benjamin Spira, Uyen Chu, Arun Narota, Hiroki Murata, Yasuaki Hashimoto, Roberto F. Nicosia, Xintao Qiu, Steve K. Huang, Maria Trojanowska, Xaralabos Varelas, Giovanni Ligresti

## Abstract

Idiopathic pulmonary fibrosis (IPF) is a progressive lung disease characterized by excessive extracellular matrix deposition and irreversible architectural distortion of the lung. Fibrotic remodeling is driven by dynamic interactions among endothelial, fibroblast, epithelial and immune cells. Although pulmonary endothelial cells (ECs) are increasingly recognized as important contributors to IPF pathogenesis, the molecular and cellular events underlying endothelial dysfunction remains poorly understood. Using integrative multi-omics analyses of human IPF lungs combined with functional in vitro assays, we identify ACKR1-expressing venous endothelial cells (ACKR1+ VECs) as critical regulators of a pathogenic niche that promotes lung fibrosis. Single-cell RNA sequencing and spatial transcriptomics analyses reveal that ACKR1+ VECs exhibit a distinct pro-fibrotic and pro-inflammatory transcriptional program enriched for hypoxia responses, extracellular matrix remodeling, and immune cell recruitment. In both mouse and human fibrotic lungs, ACKR1+ VECs localize adjacent to fibroblastic foci and are surrounded by pro-fibrotic CD68+/CCR5+/SPP1+ macrophages-monocytes, suggesting a spatial organized cellular crosstalk supporting fibrotic remodeling. Consistent with these findings, in vitro co-culture assays using ACKR1+ VECs isolated from IPF lungs demonstrate that these cells drive myeloid recruitment and fibroblast activation through ACKR1 dependent mechanisms. Silencing of ACKR1 in IPF-derived VECs suppressed inflammatory and fibrotic transcriptional programs, and pharmacological inhibition of ACKR1 attenuated stromal and immune remodeling and reduced bleomycin-induced lung fibrosis in vivo. Together, these findings identify ACKR1+ VECs as key orchestrators of fibrosis progression and establish ACKR1 and the pathogenic vasculature as promising therapeutic targets for IPF.

**Clinical Relevance:** Idiopathic pulmonary fibrosis (IPF) is a progressive and fatal lung disease with limited treatment options. We identify ACKR1-expressing venous endothelial cells as key drivers of inflammatory and fibrotic remodeling and show that pharmacologic inhibition of ACKR1 attenuates experimental lung fibrosis. These findings establish endothelial ACKR1 as a promising therapeutic target and highlight the pulmonary vasculature as a novel avenue for disease-modifying therapies in IPF.

## Introduction

Idiopathic pulmonary fibrosis (IPF) is a chronic, progressive interstitial lung disease characterized by excessive deposition of extracellular matrix (ECM), loss of alveolar architecture, and decline of pulmonary function, ultimately resulting in respiratory failure and death with a median survival period of 3-5 years^1,2^. Despite considerable advances in understanding the pathogenesis of IPF, cellular and molecular mechanisms that drive the initiation and progression of fibrosis remain poorly understood.

Pulmonary fibrosis is characterized by the establishment of a pathogenic profibrotic niche that drives the extensive deposition of collagen and other ECM proteins within the lung. Although most IPF studies have focused on epithelial and mesenchymal cells, growing evidence indicates that pulmonary vasculature dysfunction is also a key driver of fibrotic progression^3–6^. Vascular abnormalities, including increased endothelial permeability, aberrant cytokine production, and endothelial senescence have been implicated in disease progression^3,7,8^. For example, studies in aged mouse lungs with sustained fibrosis have shown that ECs show reduced expression of endothelial identity genes and increased expression of inflammatory and fibrosis-associated genes compared to young animals^3,6^. Loss of key transcription factors such as ERG and FOXF1, which regulate endothelial identity and inflammation, have been shown to increase collagen deposition, promote lung inflammation, and impair critical signaling pathways^3,9,10^. These data suggest that impairment of endothelial transcriptional programs drives a maladaptive EC state that support inflammation, disrupts vascular homeostasis, and promotes persistent fibrotic remodeling.

Pulmonary vascular remodeling is a hallmark of IPF progression and involves substantial alterations in the abundance and phenotype of distinct EC populations. Among these, aerocytes (CAP2 cells), one of the two major capillary endothelial subtypes responsible for alveolar gas exchange, are dramatically depleted within severely fibrotic regions, highlighting the disruption of specialized vascular niches during disease progression^11,12^. Conversely, we and others have shown that venous endothelial cells (VECs), defined by the expression of Atypical Chemokine Receptor 1 (ACKR1), expand within fibrotic regions^3^, suggesting selective remodeling of the pulmonary vascular bed that favors the emergence of pathogenic EC populations. This expansion highlights the remarkable plasticity of VECs and raises the possibility that they actively contribute to disease progression. Consistent with this hypothesis, VECs express adhesion molecules and chemokine receptors implicated in immune cell recruitment, including P-selectin and E-selectin, supporting a role in lung inflammatory cell trafficking and subsequent fibrotic remodeling^13,14^. Consistent with observations in human IPF lungs, we recently reported the emergence of ACKR1+ VECs in mouse lungs in response to bleomycin injury. These VECs exhibit a transcriptional program similar to their human counterparts and localize within fibrotic regions of injury, surrounded by smooth muscle actin (αSMA)-positive cells, suggesting potential paracrine communication with the cellular niche.

ACKR1 is an atypical chemokine receptor that has reported functions as a cell surface decoy receptor, regulating chemokine gradients and leukocyte trafficking ^15–17,45^. Although traditionally viewed as a scavenger receptor, emerging evidence suggests that ACKR1 can also transduce intracellular signals ^15,17,45^. Aberrant ACKR1 expression has been implicated in a range of pathological conditions, including inflammatory diseases, cancer metastasis, and vascular dysfunction^15,18–21,27^. However, whether ACKR1 contributes to lung venous remodeling or the pathogenesis of diseases, such as pulmonary fibrosis, remains unknown.

In this study, we leveraged single-cell RNA sequencing (scRNAseq) and spatial RNA sequencing from healthy and IPF lungs to dissect the transcriptional make-up and spatial distribution of these cells. We isolated ACKR1+ VECs from IPF lungs and dissected the mechanisms by which these cells drive the establishment of a profibrotic microenvironment and orchestrate intercellular communication with other cell types. We found that ACKR1+ VECs are enriched for hypoxia, glycolysis, immune recruiting and pro-inflammatory programs and localize adjacent to pro-fibrotic macrophages and scar-forming fibroblasts. In vitro co-culture experiments using isolated lung ACKR1+ VECs demonstrated that these cells establish direct contact with macrophages and act as a source of paracrine profibrotic signals that influence lung fibroblasts. Mechanistically, we identify ACKR1 as a central regulator of these pathogenic functions. ACKR1 silencing abrogates the pro-inflammatory and pro-fibrotic features of IPF-derived VECs and pharmacologic inhibition of ACKR1 suppressed the development of bleomycin-induced lung fibrosis in vivo. Collectively, these findings establish ACKR1+ VECs as direct contributors to the fibrotic niche and highlight ACKR1 signaling as a promising therapeutic target for pulmonary fibrosis.

## Results

### Expansion of ACKR1⁺ venous endothelial cells is a hallmark of progressive pulmonary fibrosis

To define EC state changes in human IPF, we analyzed a publicly available scRNA-seq dataset comprising 3 healthy and 3 IPF lungs^22^. Reclustering of all CDH5+ ECs yielded a total of 4,338 cells, comprising of 1,667 from normal lungs and 2,671 from IPF lungs (**Figure 1A**). Differential gene expression analysis comparing healthy and IPF lung revealed a distinct transcriptomic program across ECs, with markers associated with fibrotic vascular remodeling, such as *COL15A1*, and notable responders to EC stress, including ACKR1, which encodes Atypical Chemokine Receptor 1, a cell surface factor that has been previously associated with fibrotic remodeling and cancer^20^ (**Figure 1B**). Using established lineage markers, we annotated four major vascular EC subtypes from the scRNA-seq data: arterial ECs (DKK2+), general capillary (gCap) ECs (SLC6A4+), aerocyte capillary (aCap) ECs (EDNRB+), and VECs (PTGDS+)^23,24^ (**Figure 1C**, **Sup Figure 1a**). Cell composition analysis revealed a striking shift in endothelial populations in IPF lungs, characterized by an expansion of VECs accompanied by a relative reduction of arterial, gCap, and aCap ECs (**Figure 1C**). This was also evident by the loss of gCap (SLC6A4) and aCap (EDNRB) EC identity genes in the IPF lungs (**Figure 1B**). These expanded venous ECs in IPF exhibited high expression of ACKR1 (**Sup Figure 1B-D**), with a robust increase in the number of VECs expressing *ACKR1* in IPF compared to healthy lungs (**Figure 1E and Sup Figure 1C**).

**Figure 1.**
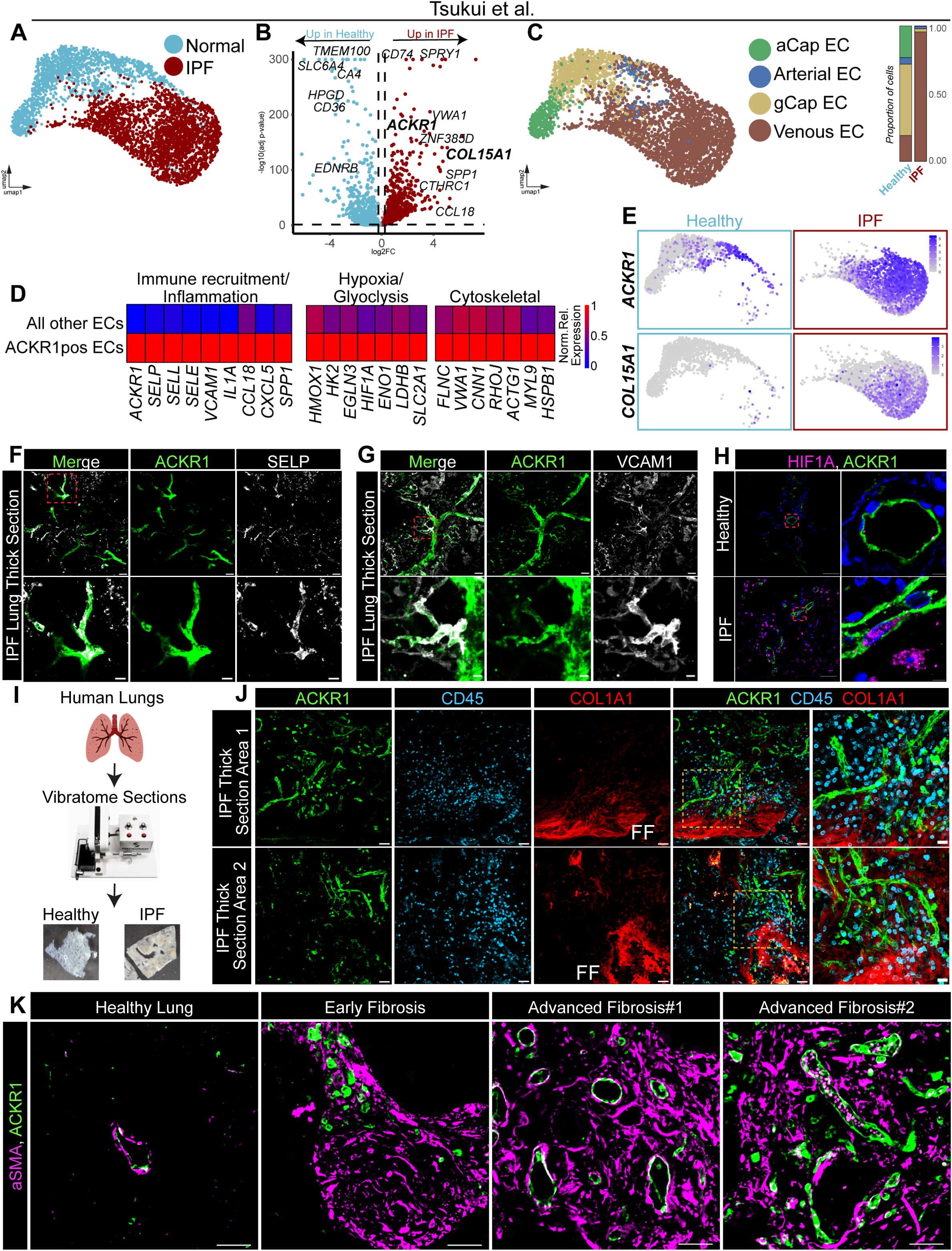
ACKR1 positive Venous Endothelial Cells are a hallmark feature of lung fibrosis. **(A)** UMAP projection of all endothelial cells from Tsukui et al.(22) (light blue-healthy red-IPF)(4,338 cells) **(B)** Volcano plot comparing genes differentially expressed across healthy and IPF endothelial cells. **(C)** UMAP projection of all endothelial cells from Tsukui et al. and proportion analysis **(D)** Heatmap of immune, hypoxic and cytoskeletal genes enriched in ACKR1pos VECs. **(E)** UMAP projection with gene expression for ACKR1 and COL15A1 split between healthy and IPF cells. **(F)** IF for ACKR1 and SELP in human IPF precision cut lung slices (large scale 50µm, small scale 20µm) **(G)** IF for ACKR1 and VCAM1 in human IPF precision cut lung slices (large scale 50µm, small scale 10µm) **(H)** IF for ACKR1 and HIF1A in healthy and IPF lungs (large scale 50µm, small scale 5µm). **(I)** Schematic for precision cut lung slices **(F)** IF for ACKR1, CD45 and COL1A1 in human IPF precision cut lung slices (large scale 50µm, small scale 20µm)(FF=Fibroblastic Foci). **(K)** IF for ACKR1 and aSMA in healthy and IPF lungs (large scale 50µm)

To gain insight into the functional properties of ACKR1+ VECs we performed gene ontology enrichment analysis on their transcriptomic programs. Compared to other ECs, ACKR1⁺ VECs were significantly enriched for pathways involved in leukocyte recruitment, cytoskeletal remodeling, and ECM remodeling (**Sup Figure 1E).** Further, differential expression of genes linked to immune recruitment and inflammation, hypoxia, glycolysis, and cytoskeletal dynamics indicates that ACKR1+ VECs adopt a distinct cellular state (**Figure1D**). To validate transcriptional signatures at the protein level and explore the spatial organization of ACKR1^+^ VECs, we performed immunohistochemistry on Precision Cut Lung Slices (PCLS) from healthy and IPF lungs (**Figure 1I).** Consistent with the scRNA-seq data, ACKR1+ VECs exhibited high levels of the immune-adhesion molecules, SELP and VCAM1 (**Figure 1F-G**). Additionally, immunostaining for HIF1A, a marker that has been shown to be associated with fibrotic vascular remodeling in IPF^25,26^, revealed elevated nuclear localization in ACKR1+ VECs (**Figure 1H).** Given that these cells are enriched for genes encoding proinflammatory and immune cell modulators, we hypothesized a role in immune recruitment. To test this, we immunostained PCLS for ACKR1, collagen-I and the pan leukocyte marker CD45. We found that ACKR1+ vessels are located in proximity to collagen dense fibrotic foci, surrounded by CD45+ immune cells (**Figure 1J**). Although ACKR1+ VECs were also detected in healthy lung tissue, both their abundance and vascular density increased markedly with fibrosis severity (**Figure 1K**). Together, these data implicate ACKR1+ VECs and associated activated phenotypes as contributors to the pro-inflammatory and pro-fibrotic environment observed in IPF.

### ACKR1+ VECs are found adjacent to scar-forming fibroblasts and pro-fibrotic macrophages in IPF lungs

To investigate the signaling relationships between ACKR1+ VECs and their surrounding microenvironment in IPF, we performed CellChat analysis to infer cell-cell communication networks from scRNA-seq data ^28^. Analysis revealed that the majority of outgoing signals from ACKR1+ VECs in IPF lungs were directed towards immune cell populations (**Figure 2A**), whereas incoming signals primarily originated from immune and mesenchymal cells (**Figure 2B**), suggesting bidirectional crosstalk between these cells. To determine whether these predicted interactions were reflected in tissue architecture, we leveraged publicly available spatial transcriptomic data from healthy and IPF lungs^29^. We generated a gene signature for ACKR1^+^ VECs from our scRNA-seq analyses and applied it to the spatial transcriptomic data. This analysis demonstrated a significant enrichment of the ACKR1+ VEC signature in IPF lung compared to healthy lungs. (**Figure 2C**). Spatial deconvolution in IPF lungs further revealed that ACKR1+ VECs are positioned in close proximity to monocyte/macrophages, mesenchymal cells, and dendritic cells (**Figure 2D**). Moreover, ACKR1+ VECs were spatially associated with both CTHRC1+ mesenchymal cells and CD14+ macrophages/monocytes, both of which are considered strong contributors to the fibrotic niche^30^(**Figure 2E&F**).

**Figure 2.**
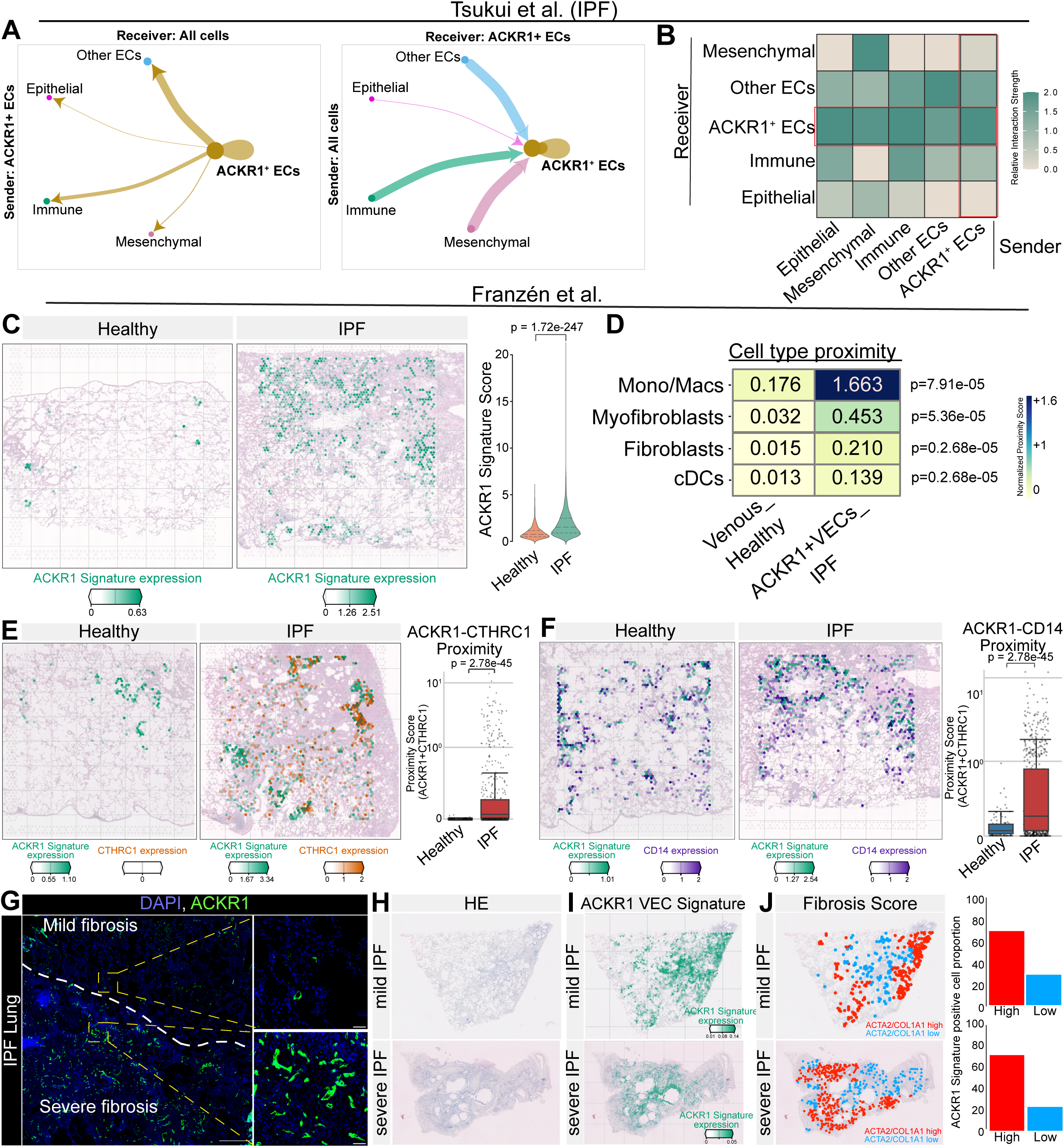
ACKR1+ VECs communicate with their microenvironment. **(A)** Circle plot showing CellChat analysis of outgoing and incoming signaling in IPF lungs and **(B)** heatmap depicting relative interaction strength between senders and receivers. (line thickness indicates relative communication probability). **(C)** Enrichment and spatial mapping of the ACKR1 VECs gene signature in human lung spatial transcriptomics data (Franzén et al.(29)). **(D)** Heatmap displaying the mean spatial proximity scores between source endothelial populations CPE+ / CDH5+ double-positive spots in Healthy Controls (left) and ACKR1 signature-scoring spots in IPF(right) and various target cell types. **(E)** Enrichment of the ACKR1 VEC-CTHRC1 gene signature across healthy and IPF samples from the *Franzén et al* (29) dataset and proximity quantification. **F)** Enrichment of the ACKR1 VEC-CD14 gene signature across healthy and IPF samples from the *Franzén et al* (29) dataset and proximity quantification. **(G)** IF for ACKR1 in an IPF lung (large scale 500µm, small scale 50µm). **(H)** HE, ACKR1 VEC signature enrichment and fibrosis score enrichment on mild and severe IPF biopsy spatial RNA. Statistical significance: **(C,E,F)** non-parametric Wilcoxon rank-sum test; **(D)** two-sided Mann–Whitney *U* test.

These findings suggested that ACKR1⁺ VECs may actively orchestrate the formation of the pro-fibrotic niche. To further investigate this possibility, we performed 10x Genomics Visium Spatial sequencing on human lungs representing distinct stages of disease, including mild IPF (early fibrosis) and severe IPF (advanced fibrosis). Each sample generated 362-446 million sequencing reads, with an average of 409 million reads per sample. Across tissue sections, we detected an average of ∼3,000 spatial transcriptomic spots per section and roughly 1,300 genes, each representing the combined transcriptome of mixture of cells in the tissue within that spot (**Sup Figure 2A-C**). Notably, severely fibrotic lungs exhibited a greater number of detected genes and higher transcript counts per spot than mildly fibrotic lungs, consistent with increased cellular density in advanced disease (**Sup Figure 2C**). Due to the abundance of ACKR1 + VECs within fibrotic regions (**Figure 2G**), we asked whether their abundance correlated with the extent of fibrosis.

Projection of the ACKR1+ VEC gene signature onto our transcriptomic datasets demonstrated enrichment in both mild and severe IPF lungs (**Figure 2H-I**). To quantify spatial relationships, we stratified spatial transcriptomic spots according to the expression of the myofibroblast markers ACTA2 and COL1A1 (ACTA2/COL1A1 high and low), classifying them as fibrosis-high or fibrosis-low regions (**Figure 2J**). Regions with high ACTA2/COL1A1 expression exhibited significantly greater enrichment of ACKR1+ VECs compared to low-expression regions, suggesting that these vessels are associated with and may contribute to the propagation of scar-forming fibroblasts across both early and advanced stages of IPF.

Due to the proximity and inferred communication with leukocytes, we revisited the CellChat analysis, which suggested that ACKR1+ VECs communicate with both macrophages and monocytes (**Figure 3A**). We found that in IPF lungs, ACKR1+ VECs have enriched signaling towards macrophages through several chemokine ligands, including CCL14 and CCL2, signifying that ACKR1+ VECs might be actively recruiting myeloid cells to the sites of fibrotic lesions (**Figure 3B**). Similarly, we found that macrophages and monocytes signal to ACKR1+ VECs through multiple proinflammatory cytokines (CCL18, CXCL8, CXCL3, CCL2) as well as ostepontin (SPP1), a key regulator of pro-fibrotic macrophage function ^31^ (**Figure 3B**). These predicted interactions were further supported by analysis of the Franzén *et al.* spatial transcriptomic dataset (**Sup Figure 2D**). To validate these observations, we performed immunostaining in IPF PCLS for CD68, CCR5 and SPP1, which are markers of macrophage/monocytes considered to contribute to fibrotic remodeling^31,32^ and demonstrate close proximity with ACKR1+ VECs (**Figure 3C-E**). We also investigated signaling interactions between mesenchymal cells and ACKR1+ VECs. CellChat analysis highlighted several ACKR1+VEC derived signals that are known to promote fibrotic responses in fibroblasts such as NAMPT, SPP1, POSTN and ANGPT2 (**Sup Figure 3B**). Conversely, myofibroblasts exhibited strong extracellular matrix- and secreted factor-related signaling toward ACKR1⁺ VECs, indicating reciprocal communication between these populations (**Sup Figure 3B**). Collectively, these findings suggest that ACKR1+ VECs actively participate in fibrotic remodeling by engaging in bidirectional crosstalk with scar-forming fibroblasts and pro-fibrotic macrophages.

**Figure 3.**
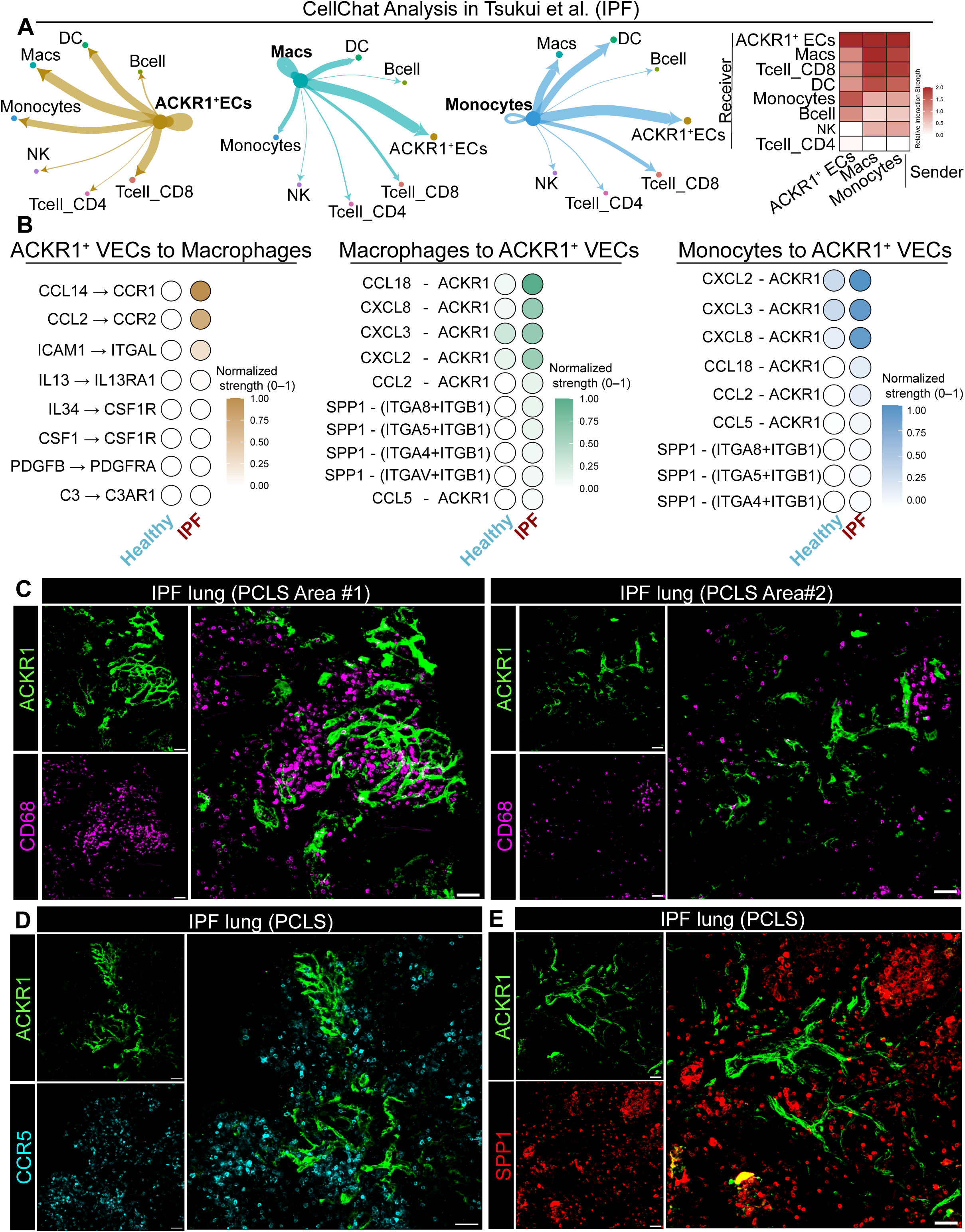
ACKR1+ VECs recruit and interact with immune cells. **(A)** Circle plot depicting CellChat analysis of outgoing and incoming signaling in IPF lungs and heatmap showing relative interaction strength between senders and receivers (line thickness indicates relative communication probability). **(B)** Bubble plot with Ligand-Receptor pair interactions between ACKR1to macrophages (left), Macrophages to ACKR1 VECs and Monocytes to ACKR1 VECs. Color indicates normalized interaction strength. **(C)** IF for ACKR1 and CD68 in IPF precision cut lung slices (scale 50µm). **(D)** IF for ACKR1 and CCR5 in IPF precision cut lung slices (scale 50µm). **(E)** IF for ACKR1 and SPP1 in IPF precision cut lung slices (scale 50µm).

### ACKR1+ VECs promote fibroblast activation and myeloid cell recruitment

To investigate the functional properties of ACKR1+ VECs, we developed a protocol to isolate and expand both ACKR1+ VECs and non-ACKR1 ECs (ACKR1-ECs) derived from the same IPF lungs. (**Figure 4A**). Both cell types retain canonical EC cobblestone morphology in vitro (**Figure 4B**), and Western blot analysis confirmed elevated ACKR1 expression in ACKR1+ VECs relative to ACKR1-ECs (**Figure 4C**). We also validated the expression of several genes that were identified in the scRNA-seq, which revealed high expression of associated inflammatory (CXCL1, CXCL5, IL18, IL1B), fibrotic (TGFB1), hypoxic (HIF1A) and metabolic (LDHA) genes^33^ (**Figure 4D**) in ACKR1+ VECs compared to non-ACKR1 ECs. Given the close spatial proximity of ACKR1+ VECs to scar-forming fibroblasts in IPF lungs, we tested whether ACKR1+ VECs have the capacity to activate lung fibroblasts in vitro. To this end, we cultured IPF-derived ACKR1+ VECs and non- ACKR1 ECs for 48 hours and then transferred the conditioned media (CM) to healthy lung fibroblasts (**Figure 4E**). Gene expression analysis revealed that conditional media derived from ACKR1 VEC conditioned media elicited the highest upregulation of CTHRC1, COL1A1 and ACTA2 when compared to ACKR1- EC, or ECs isolated from healthy lungs (**Figure 4F**). To investigate the immune-recruiting capacity of these cells, we performed an adhesion assay in which IPF-derived ACKR1+ VECs and ACKR1-ECs were cultured as monolayers, followed by the addition of GFP-expressing THP-1 monocytes for 48 hours (**Figure 4G**). This analysis demonstrated that ACKR1+ VECs exhibited significantly increased binding of THP-1 cells compared to matched ACKR1- ECs (**Figure 4H**), indicating an enhanced cell adhesion capacity. Additionally, we performed a transwell migration assay in which IPF-derived ACKR1+ VECs and ACKR1- ECs were cultured on porous membranes, followed by the addition of GFP-labeled THP- 1 cells to the apical chamber. This assay revealed increased THP-1 migration in the presence of ACKR1+ VECs compared to ACKR1-ECs. (**Figure 4I-J**). These findings demonstrate that ACKR1+ VECs function to promote both fibroblast activation and myeloid cell recruitment, supporting their role as active organizers of the pro-fibrotic vascular niche through both paracrine signaling and cell-cell interactions.

**Figure 4.**
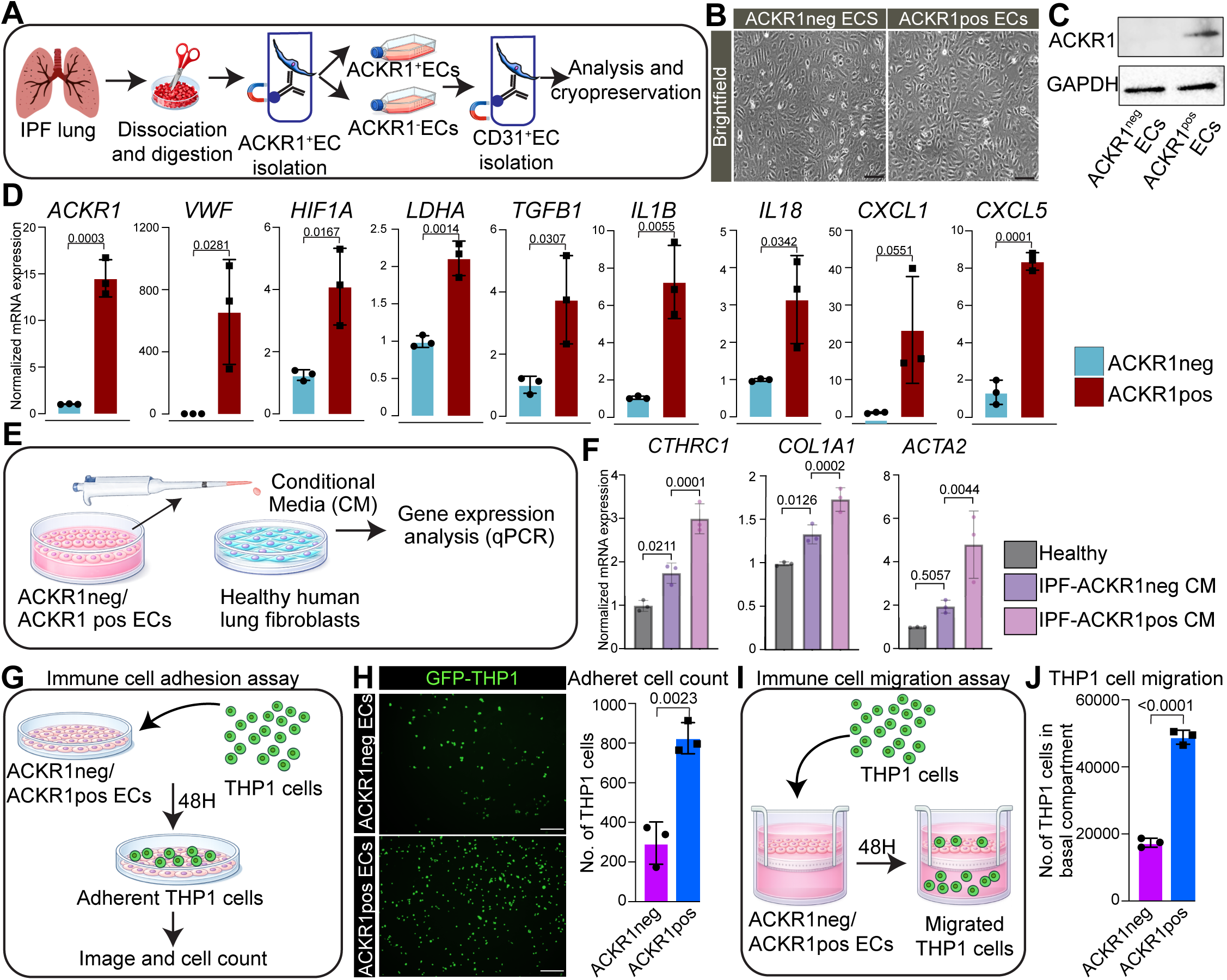
ACKR1^+^ VECs can directly remodel their microenvironment. **(A)** Schematic for ACKR1^+^ VEC isolation **(B)** Brightfield image of ACKR1^+^ and ACKR1^-^ ECs. (scale 125 pixels) **(C)** Western blot for ACKR1 **(D)** Boxplots showing normalized mRNA expression for inflammatory and hypoxic markers **(E)** Schematic for conditional media experimental set up (**F)** Boxplots showing normalized mRNA expression for CTHRC1, COL1A1 and ACTA2 **(G)** Schematic for immune cell adhesion experimental set up **(H)** Fluorescently labeled THP1 cells adhered to ACKR1^-^ and ACKR1^+^ ECs and quantification of adhered cells (scale 250 pixels). **(I)** Schematic for immune cell migration experimental set up **(J)** Quantification of migrated THP1 cells. Statistical analysis: **(D,H,J)** two-tailed Student’s t-test and **(F)** a one-way ANOVA.

### ACKR1 is required for the inflammatory and pro-fibrotic phenotype of VECs

Prior reports have suggested that ACKR1 can facilitate intracellular signaling that impact inflammatory responses^19,20^, raising the possibility that ACRK1 might actively contribute to the VEC state and pathogenic functions. To test this, we silenced ACKR1 in IPF-derived ACRK1+ VECs using siRNA, which effectively reduced ACKR1 expression compared to the scrambled control (**Figure 5A**). We then examined whether ACKR1 regulates the enhanced adhesive properties of these cells to monocytes and observed that that the number of THP1 cells interacting with the endothelial monolayer lacking ACKR1 was significantly lower than control cells with intact ACKR1 (**Figure 5B**). We next investigated whether ACKR1 contributes to the ability of VECs to activate fibroblasts. Conditioned media derived from ACKR1-silenced VECs markedly reduced fibroblast activation compared to that from control VECs, as demonstrated by the downregulation of pro-fibrotic markers, such as *COL1A1*, *FN1*, and *CTHRC1* (**Figure 5C**).

**Figure 5.**
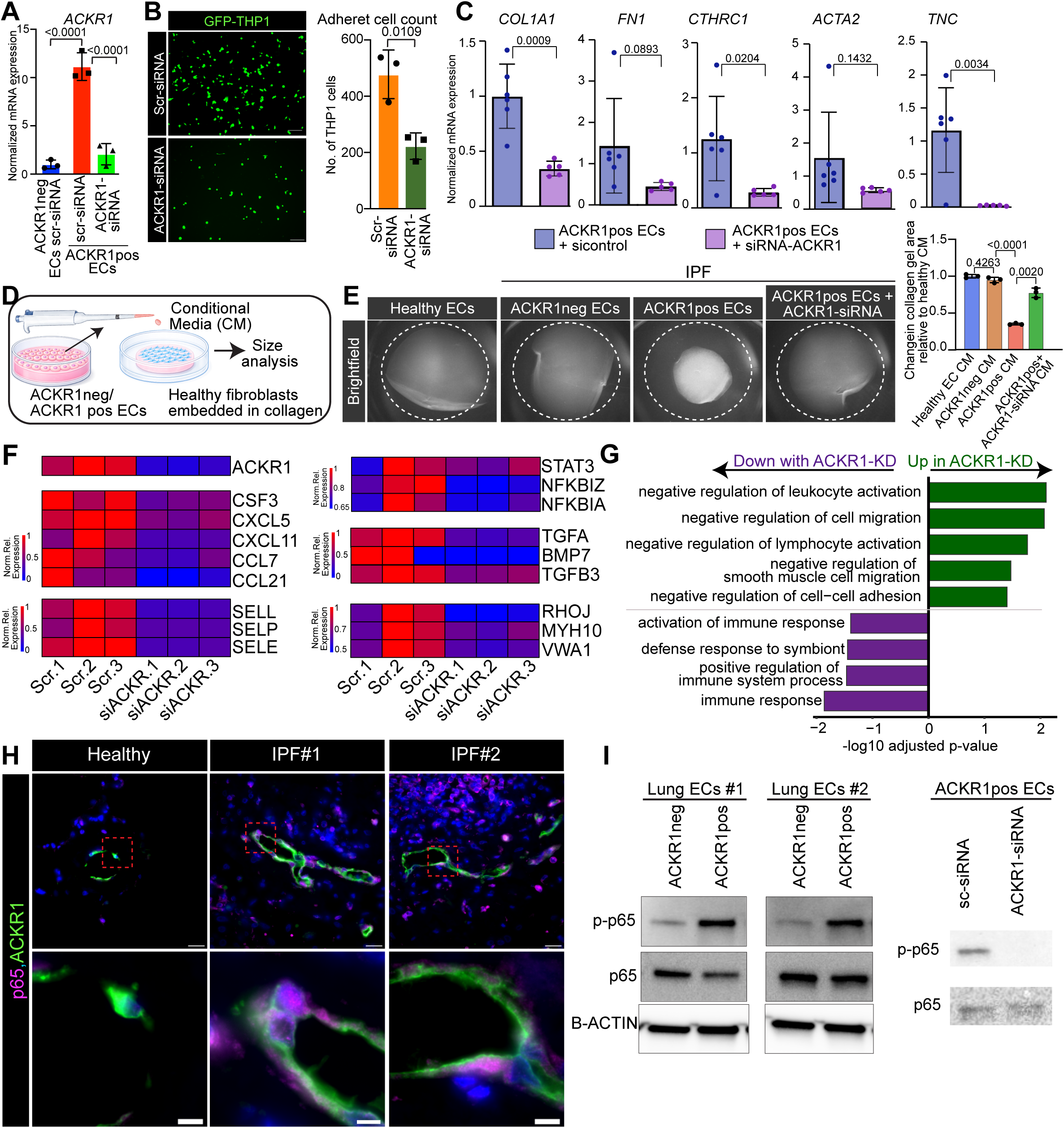
ACKR1 inhibition blocks profibrotic remodeling. **(A)** Boxplot showing normalized mRNA expression for ACKR1 **(B)** Fluorescent labeled THP1 cells adhered to Scramble or ACKR1-siRNA treated ACKR1pos ECs and quantification (scale 250 pixels) **(C)** Boxplots showing normalized mRNA expression for COL1A1, FN1, CTHRC1, ACTA2 and TNC **(D)** Schematic for collagen contraction assay experimental set up **(E)** Representative images of collagen contraction and quantification of collagen area **(F)** Heatmap for immune recruiting, profibrotic and cytoskeletal genes **(G)** GO enrichment of upregulated/downregulated genes. **(H)** IF staining for p65 and ACKR1 in healthy and IPF lungs (top scale 20µm, bottom scale 5µm). **(I)** Western blot for P-p65 and p65 Statistical significance: Statistical analysis: **(B,C)** two-tailed Student’s t-test and **(A,E)** a one-way ANOVA.

To determine whether these transcriptional changes translated into functional differences, we performed collagen gel contraction assays in which primary human lung fibroblasts were embedded in collagen matrices and treated with conditioned media from healthy ECs, IPF-derived non-ACKR1 ECs, or IPF-derived ACKR1+ VEC either treated with scrambled siRNA control or ACRK1-siRNA (**Figure 5D**). Conditioned media from ACKR1+ VECs treated with scrambled siRNA elicited the most contractile fibroblast response, as shown by the reduction in relative gel area (**Figure 5E**), whereas conditioned media from ACKR1+ VECs treated with ACKR1-siRNA evoked a far less severe contractile response in the fibroblasts (**Figure 5E**).

The marked loss of immune cell adhesion and fibroblast-activating functions in lung VECs lacking ACKR1 prompted us to investigate the molecular pathways regulated by ACKR1. To this end, we performed bulk RNA sequencing of IPF-derived ACKR1+ VECs following ACKR1 knockdown. Loss of ACKR1 resulted in the downregulation of genes encoding immune recruiting chemokines (*CXCL5, CXCL11, CCL7, CCL21),* cell adhesion molecules (*SELL, SELP, SELE*), pro-fibrotic factors (TGFA, BMP7, TGFB3), cytoskeletal proteins (*RHOJ, MYH10, VWA1*) and inflammatory responses (*STAT3, NFKBIZ, NFKBIA*) (**Figure 5F**). Gene Ontology enrichment analysis further demonstrated ACKR1 depletion suppressed inflammatory and immune-associated programs, including the negative regulation of leukocyte activation and cell-cell adhesion (**Figure 5G**). Given that the majority of upregulated factors in venous ECs lacking ACKR1 are chemokines and other inflammatory mediators, we hypothesized the involvement of NFkB, a key transcriptional regulator of inflammation^34,50^. Immunostaining for p65, a primary subunit of the NFkB pathway, in healthy and IPF lung tissues revealed strong nuclear staining in the IPF samples (**Figure 5H**). Accordingly, Western blot analysis of the activated NFkB subunit (phospho-p65) showed that ACKR1+ VECs exhibit elevated p65 phosphorylation compared to ACKR1 knockdown (**Figure 5I**). These data demonstrate that ACKR1 impacts the activation of the NF-kB signaling pathway, potentially leading to mechanisms that drive downstream transcriptional programs associated with inflammation and fibrosis.

Together, these findings identify ACKR1 as a key upstream regulator of NF-κB signaling in VECs, promoting inflammatory and pro-fibrotic transcriptional programs that drive immune cell recruitment and fibroblast activation.

### ACKR1⁺ VECs emerge early during lung injury and recapitulate the human IPF endothelial program

To better examine the dynamics behind ACKR1^+^ VEC expansion during fibrogenesis, we examined venous vessels, which are marked by the expression of Slc6a2^35^, during different stages of fibrosis development in mice following bleomycin lung injury. In sham animals, we observed very few ACKR1+ cells within venous vessels dispersed across the vasculature of the distal lung, as well as in some regions near the airways (**Figure 6A**). At day 7, 14 and 21 post bleomycin administration, however, we observed progressively increased expansion of ACKR1+ VECs (**Figure 6A**) which was accompanied by the accumulation of adjacent CD45+ immune cells (**Figure 6B**). By day 21, when fibrosis was fully established, we observed immune cell aggregates (including large numbers of macrophage/monocytes) surrounding ACKR1+ VECs, with these areas exhibiting prominent vascular sprouting and branching behavior (**Figure 6C-D**). Despite these phenotypic changes, ACKR1+ VECs maintained the expression of the venous EC marker Slc6a2, indicating preservation of venous identity (**Figure 6E**). Interestingly, venous EC sprouting has been reported in other injury contexts such as influenza^35^, suggesting that this venous-associated angiogenic program may reflect a conserved endothelial adaptation across different types of lung injury.

**Figure 6.**
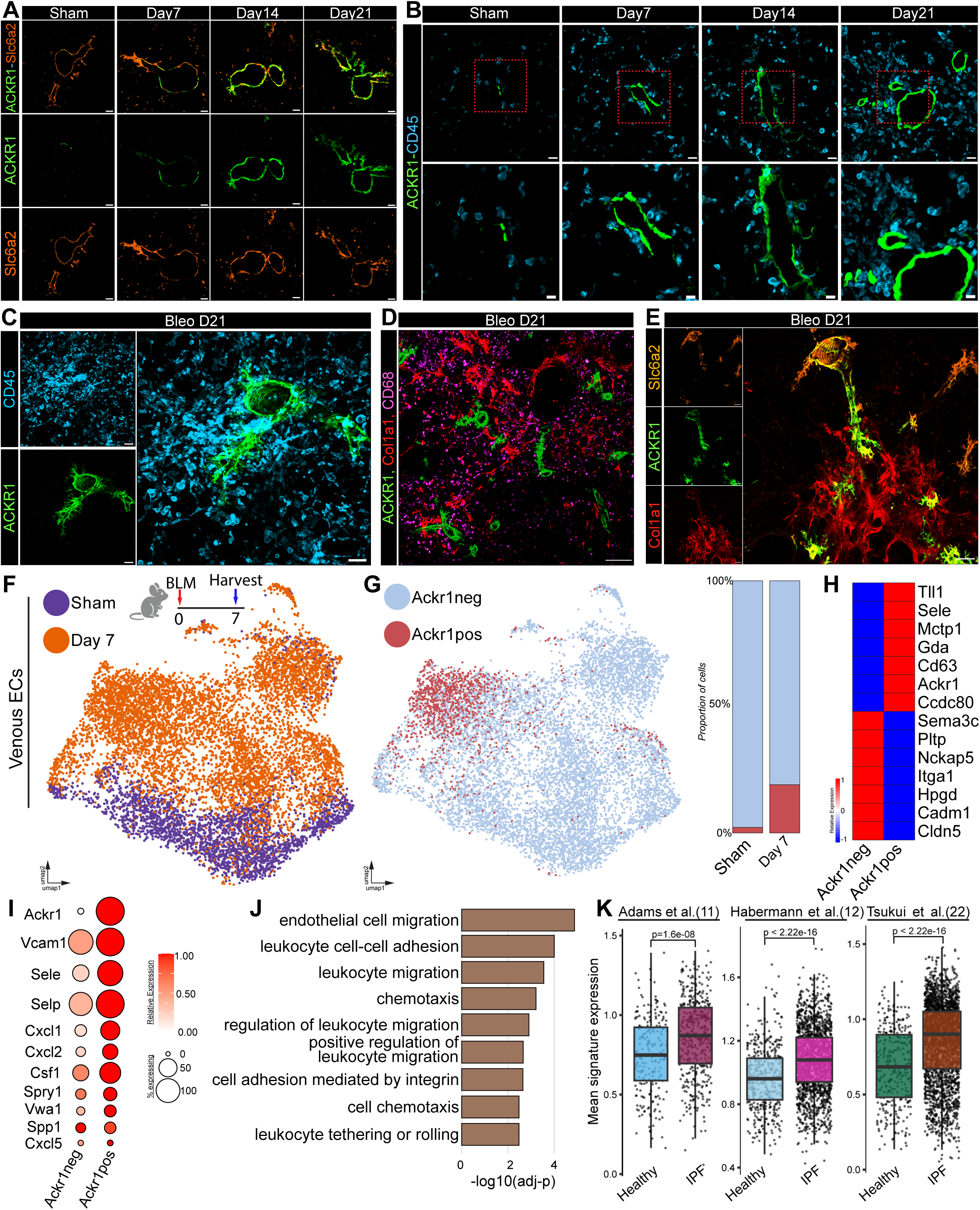
ACKR1+ VECs emergence is an early event during murine fibrosis. **(A)** IF for ACKR1 and Slc6a2 in bleomycin treated mouse lungs (scale 20 µm). **(B)** IF for ACKR1 and CD45 in bleomycin treated mouse lungs (large scale 20µm, small scale 10µm). **(C)** IF for ACKR1 and CD45 in mouse lungs twenty-one days after bleomycin installation (scale 50µm). **(D)** IF for ACKR1, CD68 and Col1a1 in mouse lungs twenty-one days after bleomycin installation (scale 200 µm). **(E)** IF for ACKR1, Slc6a2 and Col1a1 in mouse lungs twenty-one days after bleomycin installation (scale 100 µm). **(F)** scRNAseq set-up and UMAP projection of all venous endothelial cells (purple-sham, orange-seven days post bleomycin administration (cells) **(G)** UMAP projection of ACKR1 positive and ACKR1 negative cells and proportion plot **(H)** Heatmap with differentially expressed genes. **(I)** Bubble plot with inflammatory and immune recruiting genes **(G)** GO enrichment of upregulated genes. **(K)** Day seven ACKR1 signature on the Adams et al(11)., Habermann et al.(12) and Tsukui et al. (22) datasets. Statistical significance was evaluated using a two-sided Wilcoxon rank-sum test.

To dissect early molecular events that accompany the emergence of ACKR1+ VEC during the onset of lung fibrosis, we performed scRNA-seq on sham and day 7 bleomycin-injured mouse lungs. We identified 34,315 endothelial cells (12,313 from sham and 22,002 from injured lungs), which were annotated using established endothelial lineage markers (**Sup Figure 6A-C**). Although we found that many cell clusters showed transcriptional differences in response to injury (**Sup Figure 4D**), we focused our analysis on venous ECs (9914 cells; 2780 sham and 7134 injured) (**Figure 6F).**

Reannotation based on Ackr1 expression identified 1,424 ACKR1+ and 8,490 non- ACKR1 VECs (**Figure 6G**). Approximately 20% of VECs expressed ACKR1 by day 7 after injury, whereas only a small fraction were ACKR1+ VECs under homeostatic conditions, consistent with our earlier observations (**Figure 6G**). Amongst the most highly upregulated genes in the ACKR1+ VECs were *Sele* and *Ackr1* **(Figure 6H)**, while these cells also expressed several inflammatory (*Cxcl1, Cxcl2, Cxcl5, Csf1*) and immune recruiting (*Vcam1, Sele, Selp*) genes similarly to human ACKR1+ VECs (**Figure 6I and Sup Figure 4E**). Gene ontology analysis of genes associated with ACKR1+ VECs revealed enrichment of pathways linked to leukocyte recruitment and cell adhesion (**Figure 6J**), while KEGG analysis identified activation of extracellular matrix remodeling and metabolic pathways (**Sup Figure 4G**). To examine whether activated mouse ACKR1+ VECs share transcriptional features with their human counterparts, we generated a gene signature score derived from day 7 mouse ACKR1+ VECs and projected it onto three independent human scRNA-seq datasets from healthy and IPF human lungs (**Figure 6K**). This analysis revealed a robust enrichment of the mouse ACKR1+ VEC signature in IPF ECs, with minimal enrichment in healthy controls (**Sup Figure 4F**). Together, these findings indicate that the mouse model faithfully recapitulates key aspects of ACKR1+ VEC remodeling observed in human IPF lungs.

While our data suggest that injury elevates Ackr1 expression in lung venous ECs, we cannot exclude the possibility that increased cellular proliferation may also contribute to the expansion of these cells. To test this, we injured mouse lungs with bleomycin, treated mice with Edu, and isolated lungs at day 3 post injury to assess proliferation. Strong EdU+ staining was observed in ACKR1+ VECs **(Sup Figure 4I)**, suggesting that cell proliferation may in part be responsible for the accumulation of these cells under injury conditions. We also observed pronounced perivascular proliferation in cells surrounding ACKR1+ VECs, supporting the existence of a venous niche that may contribute to the onset of fibrotic development **(Sup Figure 4I)**.

### Pharmacological inhibition of ACKR1 attenuates pulmonary fibrosis

Our in vivo and in vitro data demonstrate that Ackr1 expression is increased in lung venous ECs following bleomycin challenge, and that Ackr1 silencing in human lung VECs attenuates pro-inflammatory and pro-fibrotic responses. These findings support a functional role for Ackr1 in driving injury-induced inflammatory and fibrotic activation within the venous endothelial compartment. To test this hypothesis and translate our findings to fibrotic phenotypes in vivo, we employed bleomycin-injury and tested whether pharmacological inhibition of ACKR1 attenuates fibrotic responses. To this end, we used amikacin, an antibiotic recently reported to bind and inhibit ACKR1 function^19^. Amikacin was administered daily after bleomycin injury, and lungs were harvested 21 days following bleomycin injury (**Figure 7A**).

**Figure 7.**
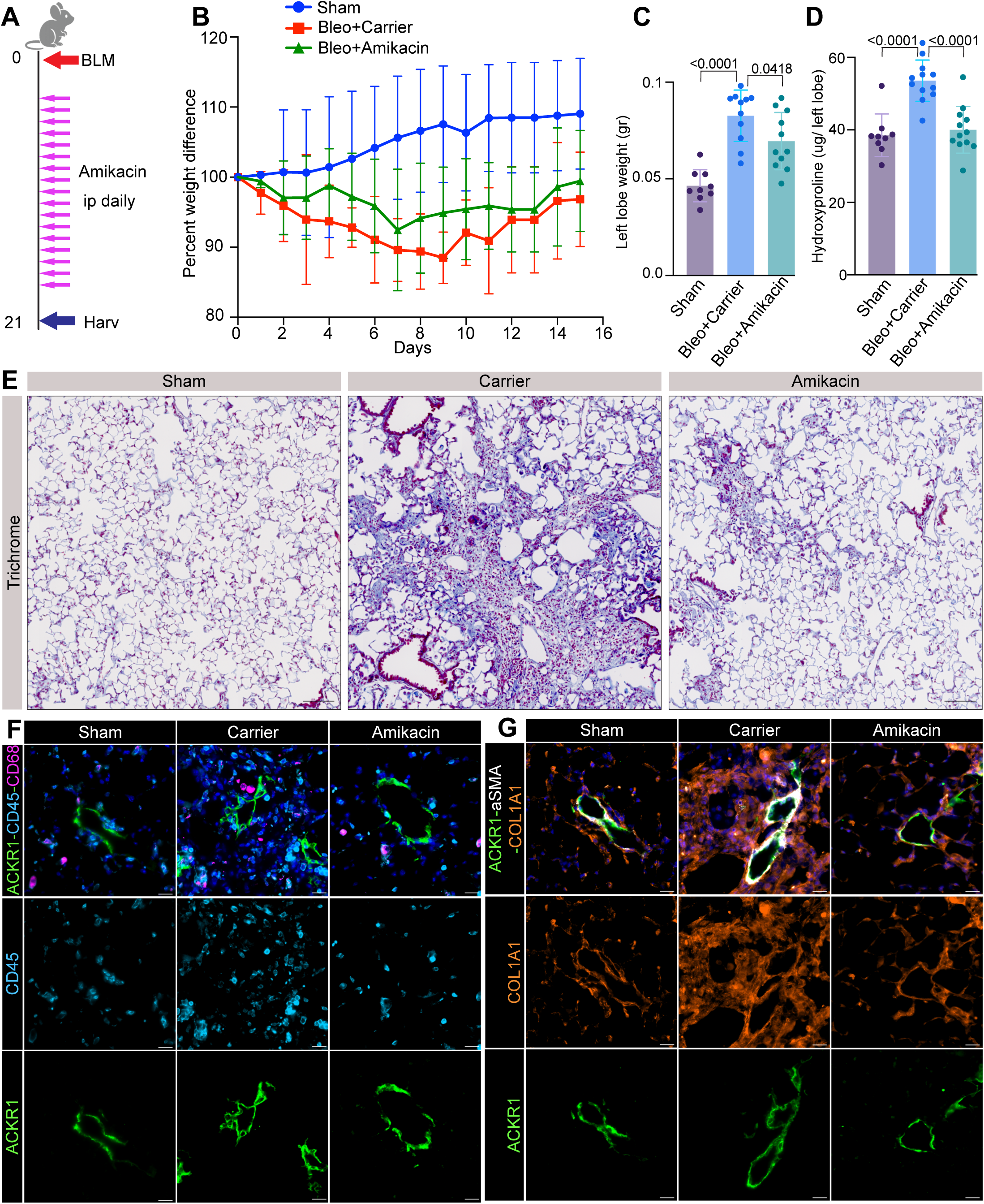
ACKR1 receptor inhibition ameliorates fibrosis in mice. **(A)** Experimental set up. **(B)** Diagram with representative body weight. **(C)** Bar plot depicting left lobe wet weight. **(D)** Bar plot depicting left lobe hydroxyproline measurement **(E)** Masson’s trichrome staining of sham, carrier and amikacin treated lungs (scale 100 µm). **(F)** IF for ACKR1, CD45 and CD68 in sham, carrier and amikacin treated lungs (scale 20 µm). **(G)** IF for ACKR1, aSMA and Col1a1 in sham, carrier and amikacin treated lungs (scale 20 µm). Statistical significance: **(C,D)** one-way ANOVA.

Pharmacological inhibition of ACKR1 markedly reduced fibrosis severity, as demonstrated by decreased body weight loss, reduced left lung wet weight, and significantly lower collagen accumulation quantified by hydroxyproline assay (**Figure 7B-D**). Consistent with these findings, Masson’s trichrome staining revealed reduced fibrotic remodeling in amikacin-treated mice (**Figure 7E**). Immunostaining for CD68, CD45, and collagen I further demonstrated diminished inflammatory cell accumulation and extracellular matrix deposition following ACKR1 inhibition (**Figure 7F,G**). Collectively, these findings demonstrate that pharmacological inhibition of ACKR1 mitigates lung inflammation and fibrosis in vivo, supporting ACKR1 as a promising therapeutic target for idiopathic pulmonary fibrosis.

## Discussion

IPF is a progressive, irreversible, and highly fatal lung disease with a median survival of 3-5 years and limited therapeutic options^36^. Although the pulmonary vasculature constitutes more than one-third of all cells in a healthy lung, the contributions of ECs in the pathogenesis of IPF is not well understood. Growing evidence indicates that the ECs in IPF are phenotypically abnormal and functionally impaired^3,4,11,37,38^, exhibiting reduced barrier integrity, aberrant inflammatory signaling, and heightened responsiveness to fibrotic stimuli^39–41^. Notably, remodeling of pulmonary venous vessels appears to be a particularly early event in IPF pathogenesis, with our work and others suggesting that VECs acquire inflammatory, angiogenic and immune cell recruiting properties during disease progression ^4,3,38^

In this study, we identify ACKR1+ VECs as key regulators of immune-stromal crosstalk and driver of fibrotic progression in IPF. By integrating scRNA-seq, spatial transcriptomics, and functional in vitro and in vivo approaches, we demonstrate that ACKR1+ VECs expand during fibrotic remodeling in both human IPF and experimental lung fibrosis and actively promote the establishment of a pro-fibrotic vascular niche. ACKR1⁺ VECs exhibit a distinct activated phenotype characterized by enhanced expression of genes associated with endothelial activation, leukocyte recruitment, hypoxia, glycolytic metabolism, and extracellular matrix remodeling. Functionally, these cells promote fibroblast activation and myeloid cell recruitment, while genetic and pharmacological inhibition of ACKR1 attenuates these pathogenic responses, establishing ACKR1 as a critical regulator of the venous endothelial phenotype. These findings build upon and substantially extend our previous work demonstrating that VECs emerging after bleomycin-induced lung injury acquire a distinct transcriptional state marked by ACKR1 expression ^3^, providing mechanistic evidence that this endothelial population actively contributes to the initiation and progression of the fibrotic niche.

ACKR1 is an atypical chemokine receptor capable of binding a broad range of inflammatory chemokines, including members of the CXC and CC chemokine families. Unlike conventional chemokine receptors, ACKR1 lacks the canonical Gαi-coupling motif required for classical G protein signaling and has therefore traditionally been viewed as a decoy or scavenger receptor that regulates chemokine availability across the endothelial barrier^18,43,44^. By shaping chemokine gradients, ACKR1 has been proposed to facilitate leukocyte adhesion, extravasation, and tissue infiltration, placing this receptor at the center of inflammatory cell trafficking^43–45^. However, increasing evidence suggests that ACKR1 also mediates non-canonical intracellular signaling^46^. Consistent with this signaling premise, we found extensive communication between ACKR1+ VECs and immune cells in IPF lungs. ACKR1+ VECs expressed high levels of secreted and adhesion molecules responsible for immune cell recruitment such as *SELP*, *SELE, VCAM1, CXCL1, CXCL2, and CSF1*^46^. Furthermore, the expansion of ACKR1+ vessels in IPF, was coupled with their close spatial association to pro-fibrotic fibroblasts and infiltrating immune cells. CellChat analysis predicted enhanced crosstalk between ACKR1+ VECs and macrophages mediated through CXCL chemokines and SPP1 signaling. Spatial transcriptomics further suggested that ACKR1+ VECs are predominantly found in proximity with SPP1+ and CCR5+ macrophages, a well characterized pathogenic macrophage population in IPF^31,32^. Notably, similar spatial associations between ACKR1+ VECs and profibrotic macrophages have been reported in other fibrotic conditions, including scleroderma, as well as in vascular pathologies such as thoracic aortic aneurysm and dissection (TAAD)^18,3,44,46^. In line with these observations, ACKR1+ VECs have been shown to directly interact with proinflammatory IL1B+ macrophages^49^, supporting a conserved role for these venous ECs in orchestrating immune cell recruitment and activation across diverse disease contexts.

Our findings demonstrate that ACKR1 is not merely a marker of venous endothelial identity but is functionally required for the pathogenic phenotype of associated with these cells. Silencing ACKR1 expression in IPF-derived VECs markedly impaired monocyte adhesion and recruitment while reducing the ability of endothelial-derived conditioned media to activate lung fibroblasts. These functional changes were accompanied by broad transcriptional reprogramming, including suppression of inflammatory chemokines, adhesion molecules, extracellular matrix-associated genes, and pro-fibrotic signaling pathways. Notably, ACKR1 knockdown reduced NFkB activation, suggesting that ACKR1 promotes a sustained inflammatory endothelial state through non-canonical signaling mechanisms that converge on NFkB-dependent transcriptional programs. Emerging evidence from other vascular diseases also supports a functional intracellular role for ACKR1. In thoracic aortic aneurysm and dissection (TAAD), ACKR1-expressing ECs regulate macrophage recruitment and inflammatory polarization through an ACKR1- NFkB-SPP1 signaling axis, and pharmacological inhibition of ACKR1 with amikacin attenuates disease progression in experimental models^19^. These observations closely parallel our findings in lung fibrosis development, where genetic silencing of ACKR1 reduced endothelial pathogenicity, and pharmacological inhibition of ACKR1 attenuated bleomycin-induced lung fibrosis. Together, these studies support the existence of a conserved ACKR1+ endothelial activation program that promotes tissue remodeling across multiple chronic inflammatory diseases. Targeting ACKR1 signaling or selectively disrupting the pathogenic functions of ACKR1+ VECs, may therefore represent a promising therapeutic strategy to limit vascular-driven inflammation and fibrotic progression in IPF.

## Material and Methods

### Ethical statement

Lung specimens from IPF patients were collected from explanted lungs at the time of transplantation. Written informed consent was obtained from all patients, and the study received approval from the University of Michigan Institutional Review Board, Ann Arbor, MI (HUM00105694). IPF diagnoses were determined using clinical pathological criteria and verified through multidisciplinary consensus conference. Supplementary IPF lung tissue was acquired from a deceased subject; next of kin authorized the autopsy in this instance (VA Medical Center, Seattle, WA). Healthy control lungs were sourced from deceased donors whose lungs were rejected for transplantation, supplied by Gift of Life, Michigan, with family consent permitting research use of the tissue. Neither patients providing IPF samples nor families donating normal control lungs received any compensation. Sex and/or gender was not factored into the study design. Both sexes participated in the study and were randomly distributed across experiments, with no sex-related differences detected in our results. Given the small number of patient specimens analyzed, statistical correction for sex or gender analysis was not conducted. This study utilized lung tissues from three healthy donors (one female and two males) along with three IPF patients (one female and two males). All animal experiments were performed in accordance with protocols sanctioned by the Institutional Animal Care and Use Committee (IACUC) at Boston University (Protocol # PROTO201900054) and adhering to the Animal Research: Reporting of In Vivo Experiments (ARRIVE) guidelines.

### Mice

All animal procedures were carried out following protocols sanctioned by the Institutional Animal Care and Use Committee (IACUC) at Boston University and adhering to the Animal Research: Reporting of In Vivo Experiments (ARRIVE) guidelines. C57BL/6J mice (Jax strain 000664) were used for these studies. All mice were given unrestricted access to food and water under a 12 h/12 h light/dark cycle, at an ambient temperature of 77-78 °F and humidity of 46–49%.

### Bleomycin-induced lung fibrosis model

Pulmonary fibrosis was triggered in 8-14 week-old male and female mice through oropharyngeal delivery of bleomycin sulfate (1.5 U/kg; Fresenius Kabi, Lake Zurich, IL, USA) dissolved in 50 µL sterile PBS, following previously published methods². Briefly, mice were anesthetized with isoflurane, and once light plane anesthesia was reached, the animals were positioned on a mouse intubation platform. The mouse’s tongue was extended and held with forceps, bleomycin was delivered into the distal portion of the oral cavity (oro-pharynx) via pipette, and the nose was lightly closed until the liquid passed down the respiratory tract. The tongue was released until gasping fully ceased, respiration was carefully observed, and the animals were positioned vertically. After recovering from anesthesia, animals were returned to their cages. Sham control mice were administered PBS. Body weight was tracked daily. At lung harvesting, mice were euthanized using 100 µl of FATAL-PLUS solution (Vortech pharmaceuticals, Dearborn, MI, USA). Hydroxyproline measurements were conducted 20 days following the final tamoxifen dose by perfusing, harvesting, and homogenizing the left lobe with a TissueLyser 2 (Qiagen) and applying a commercially available kit to quantify hydroxyproline content (Sigma MAK569). Hydroxyproline content is reported as micrograms of collagen per milligram of left lobe homogenate (μg/mg).

### ACKR1 Cell Isolation

Human lung tissue was first minced with a pair of scissors, then incubated in freshly prepared digestion buffer: Endothelial Cell Growth Medium MV 2 with supplements (PromoCell, C-22121), Antimycotic(Fisher, 15-240-096), sodium pyruvate (Fisher, 11360070), MEM non-essential amino acids (Fisher, 11140050), collagenase II (Fisher, Cat. No. 17101015), collagenase IV (Worthington, LS004188), and DNase I (Qiagen, 79254). Next, approximately 1-2 ml of minced tissue was transferred into each gentleMACS C Tube (Miltenyi Biotec, Cat. No. 130-096-334), followed by 5 ml of digestion buffer. Samples were mechanically dissociated using the gentleMACS Dissociator (Miltenyi Biotec, Order No. 130-093-235). After dissociation, samples were incubated at 37°C for 1 hour on a rotator. The digested tissue suspension was then filtered twice through 100 µm filters, and twice through 40µm filters followed by ACK lysis buffer for 5 minutes to lyse erythrocytes. The reaction was neutralized with endothelial cell medium, and the cells were centrifuged again at 1000 rpm for 5 minutes. If the pellet remained visibly red, ACK lysis was repeated. The final cell pellet was resuspended in 200µL of DMEM (Fisher, 11965092) and 4µL of DARC-PE (Biotechne, FAB4139P) for 30 minutes at 4C, washed three times, incubated with Anti-PE magnetically conjugated beads (Miltenyi, 130-048-801) and finally passed through a magnetic column(Miltenyi, 130-042-401). The cells adhered to the column were then plated on a 6 well plate, while the cells in the flow-through were plated on a T75 flask (Fisher, 12565349), and were cultured in Endothelial Cell Growth Medium MV 2 with supplements (PromoCell, C-22121). Both fractions were allowed to grow till confluency, and were then dissociated, pelleted, resuspended in 200µL of DMEM (Fisher, 11965092) and 4µL of anti-CD31 magnetically conjugated (Miltenyi, 130-091-935) and passed through a magnetic column to remove any contaminant cells.

### Cell culture

zsGreen-THP1 cells AATC (TIB202), were maintained in RPMI (Fisher, 11875101), 10% FBS (Fisher, A5670101) and Penicillin-Streptomycin (Fisher, 15140163). Human healthy fibroblasts were maintained on plastic plates in in DMEM (Fisher, 11995065), 10% FBS (Fisher, A5670101) and Penicillin-Streptomycin (Fisher, 15140163). ACKR1+ VECs were grown to confluency and then treated with then treated with ACKR1 or scramble siRNA (Horizon Discovery/Dharmacon L-L-003865-00-0020) for 48hrs.

### In vitro migration and adhesion assays

ACKR1+ VECs were cultured in a six well plate (adhesion assay) or a 0.4µm transwell (migration assay)(Greinger till they reached confluency. Then 100.000 zsGreen-THP1 cells were added to the apical side and were allowed 48 hours to adhere or migrate.

### Conditional media assay

ACKR1+ VECs were grown to confluency and then the conditioned media was harvested, spun down to remove any cells and then applied on healthy lung fibroblasts (cat) for 48 hours. The fibroblasts were then lysed and RNA extraction was performed.

### Immunohistochemistry

Formalin-fixed paraffin-embedded (FFPE) mouse lungs were sectioned serially (5 μm). Sections underwent deparaffinization following a standardized protocol and were blocked with 10% donkey serum (Sigma, S30-100ML) along with rodent antibody blocking solution (Biocare Medical, RBM961L). Staining involved primary antibodies diluted in 5% donkey serum and incubated overnight at 4°C, followed by washing and incubation with secondary antibodies and DAPI for one hour. Antibodies are detailed in Table SX. Additional stains employed the following reagents: Shandon Harris Hematoxylin (Fisher 6765003) and Trichrome stain (Masson) (Sigma HT15-1KT). Slides were dehydrated and mounted with FluorSave (Sigma 345789-20ML). Actin staining of fixed cells used Alex488-Phalloidin (Cytoskeleton SKU:PHDG1). Imaging was conducted on a Zeiss Axio Scan.Z1 microscope with Zeiss ZEN 3.1 Blue software or a Zeiss Axio D1 observer inverted microscope with Zeiss ZEN 3.1 Blue software. Image analysis was completed using CellProfiler (4.2.8) and EBImage (4.46.0).

### Bulk RNA Sequencing and gene pathway analysis

IPF-derived ACKR1+ VEC cells were cultured and treated with either scramble siRNA (Horizon Discovery/Dharmacon, D-001810-01-05) or siRNA targeting ACKR1 (MedChemExpress, 5500343490) for two days in OptiMEM media (Fisher, 31985070) for 48 hours, after which the cells were lysed and RNA extracted.Total RNA integrity was verified using RNA 6000 Pico Assay run on an Agilent 2100 Bioanalyzer (Agilent Technologies, CA, USA). Libraries were prepared using Illumina’s Stranded mRNA Library Prep, Ligation kit according to the manufacturer’s protocol (Illumina, CA, USA). To enrich for mRNA, 100 ng of total RNA for each sample was incubated with magnetic Oligo(dT) beads. Enriched mRNA was fragmented at 94C for 8 minutes. First strand cDNA was generated, and incorporation of dUTP occurred during synthesis of second strand cDNA to indicate strand specificity. Double-stranded cDNA then underwent A-tailing and ligation of pre-index anchors. Adapters and sample-specific unique dual indices were incorporated during PCR amplification (13 cycles) according to the manufacturer’s protocol (Illumina, CA, USA). Size distribution and molarity of amplified cDNA libraries were assessed via Bioanalyzer DNA 1000 Assay (Agilent Technologies, CA, USA). All libraries were sequenced on an Illumina NextSeq 2000 instrument with 550 pM input and 2% PhiX (Illumina, CA, USA).

### Quantitative Real-time PCR

Total mRNA was extracted using Quick-RNATM Miniprep (Zymo Research,Z5223), with concentration and purity assessed via Nanodrop. cDNA synthesis was performed using the High-capacity cDNA Reverse Transcription Kit (Applied Biosystems, 4368814); RT-PCR was conducted with PowerUp™ SYBR™ Green Master Mix (Applied Biosystems, A25743) and analyzed on a Step-One-Plus Real-Time PCR system (Applied Biosystems). Primer sequences employed in this study are presented in SupTable1.

### Protein extraction and Western blotting

Cells were rinsed twice with ice-cold 1x PBS, then lysed using M-PER (Fisher, 78501) supplemented with Halt Protease and Phosphatase Inhibitor Cocktail (Fisher, 78442). Lysates underwent centrifugation, and supernatants were retained. Protein concentration was determined by BSA assay kit (Fisher, A55865). Approximately 15-20 µg of protein was loaded per lane on a Mini-PROTEAN TGX 4-15% Gel (Bio-Rad, 4561084), resolved by electrophoresis, and transferred to PVDF membranes. Membranes were subsequently blocked with 5% non-fat dry milk or 5% BSA in 1x TBST (Bio-Rad Laboratories) and incubated overnight at 4°C with primary antibodies. Blots were rinsed with 1x TBST and incubated with appropriate secondary antibodies for 1 hour at room temperature. Bands were detected using Super Signal West Pico Plus (Fisher, 34580) and a ChemiDoc Imaging System (Bio-Rad), per manufacturer’s instructions.

### Spatial RNA data generation

Lung biopsies from two patients were processed for spatial transcriptomic profiling using the Visium CytAssist Spatial Gene Expression workflow according to the manufacturer’s instructions (10x Genomics, CG000495). Briefly, FFPE sections mounted on standard glass slides were deparaffinized, stained, and imaged to preserve tissue morphology for downstream spatial registration. Sections were then processed through decrosslinking and whole-transcriptome probe hybridization using the appropriate species-specific Visium CytAssist probe set. Following probe hybridization and ligation, the tissue slide was loaded onto the Visium CytAssist instrument together with a Visium CytAssist Spatial Gene Expression slide containing spatially barcoded capture areas. Ligated probes were transferred from the tissue section to the barcoded slide, thereby preserving spatial information across the tissue.

Captured probes were subsequently extended, amplified, and converted into dual-indexed sequencing libraries following the 10x Genomics library construction workflow. Final libraries were assessed for quality and concentration, pooled, and sequenced for spatial gene expression analysis. Spatially barcoded sequencing reads were later aligned with the corresponding tissue image to generate spot-level gene expression matrices. Source:

### Single-cell RNA data generation

Lungs from 8–14-week-old male mice (3 sham and 3 day-seven post-bleomycin) were harvested and dissociated into single-cell suspensions using established protocols. Briefly, mice were euthanized and perfused through the right ventricle with 15 mL cold PBS. Lungs were then inflated with DMEM containing 0.2 mg/mL collagenase and 100 U/mL DNase I, incubated for 2 minutes at room temperature, removed, finely minced, and digested in an additional 30 mL of enzyme solution for 35 minutes at 37°C with rotation. Digestion was stopped by adding DMEM supplemented with 10% fetal bovine serum, and the resulting suspension was passed through a 40-µm strainer to eliminate debris. Red blood cells were lysed by resuspending the pellet in 1.5 mL lysis buffer for 90 seconds, followed by dilution with 9 mL PBS. The homogenate was incubated with anti-CD45 microbeads (Miltenyi 130110618) for 30 minutes at 4°C and then passed through an isolation column (Miltenyi 130042401). Cells were washed and resuspended in 0.2 mL 1× PBS containing 0.04% BSA before submission to the Single-Cell Sequencing Core. Cell number and viability were assessed using a manual hemocytometer.

Single cell suspensions were then processed using the Chromium Single Cell 3’ Reagent Kit v4 according to the manufacturer protocol (10x Genomics, CA, USA). Cell suspensions were loaded on a Chromium GEM-X Single Cell 3’ Chip at concentrations ranging between 780,000 - 1,910,000 cells/mL with viability from 84-91% for a targeted cell recovery of 20,000 cells per sample. The size distribution and molarity of amplified libraries were assessed using the Bioanalyzer High Sensitivity DNA Assay (Agilent Technologies, CA, USA). The libraries were then pooled and sequenced twice on a P4 100 cycle flow cell and once on a P3 100 cycle flow cell via the Illumina NextSeq 2000 instrument (Illumina, CA, USA) resulting in 123,901,682 total RNA counts. There were 34315 cells captured with an average of 1517.2 genes per cell.

### Preprocessing of spatial Visium RNA-seq data

Data was retrieved from https://www.ebi.ac.uk/biostudies/studies/S-BSST1410, as described in Franzén et al. (90), with the article and data made available under a Creative Commons Attribution 4.0 International License (http://creativecommons.org/licenses/by/4.0/). The cell2location annotation was sourced from ’hs_visium_stutility_obj.rds’. SCANPY (version 1.11.5) and Squidpy (1.6.5) were used for preprocessing. The Space Ranger output files together with the corresponding histology images were combined into a single anndata object. Quality control parameters were evaluated, and the data were filtered to exclude spots with fewer than 800 counts, more than 45000 counts, fewer than 800 genes, or fewer than 25 cells. To filter Visium spatial data, we implemented a quality control pipeline with Squidpy to segment and count cellular units inside each capture spot. High-resolution H&E images were initially preprocessed with a Gaussian smooth (sigma = 1) and segmented via a Watershed algorithm to delineate individual cellular boundaries. By scaling the transcriptomic coordinates to image pixel space, we extracted the number of distinct segmentation labels underneath each spot. Finally, we imposed a stringent filtering threshold, keeping only those spots with more than 30 segments to guarantee adequate cellular density for reliable downstream analysis and deconvolution.

### Spatial Proximity Analysis

To measure the spatial relationship between specific cell populations and ACKR1+ VECs niches, we computed a Spatial Proximity Score P for each Visium tissue patch. This metric considers the physical adjacency of spots on the hexagonal grid rather than basic global correlation.

### Graph Construction

For each tissue section, a spatial adjacency graph was generated using the Squidpy package. Spots were defined as neighbors *W_ij_* = 1 if they shared a direct border on the Visium grid; otherwise, *W_ij_* = 0.

### Proximity Score Calculation

Spatial proximity between two variables, *A* (e.g., ACKR1 gene signature score) and *B* (e.g., cell-type abundance or gene signature score), was quantified using the following metric:

Where:

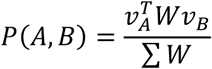

*v_A_* and *v_B_* are vectors containing the normalized expression or abundance values across all spots within a tissue section. The numerator quantifies the cumulative interaction between neighboring spots, while the denominator corresponds to the total number of edges in the spatial graph, thereby normalizing the score for differences in tissue size and graph connectivity.

Statistical significance of proximity scores between groups was assessed using the non-parametric Wilcoxon rank-sum (Mann–Whitney U) test.

### Spatial Visualization

Spatial distributions of gene signatures and predicted cell-type abundances were visualized using the cell2location and Scanpy packages. Raw expression values or cell2location abundance estimates were first extracted for each sample and subsequently normalized. These values were then overlaid onto high-resolution H&E images to generate spatial expression maps. For comparisons between healthy and IPF tissues, CPE/CDH5-positive spots were annotated as healthy venous endothelial cells, while the ACKR1 gene signature was used to identify regenerative ACKR1 VECs. The spatial proximity analysis described above was then performed to quantify the association between ACKR1+ and neighboring cell populations, with resulting proximity scores displayed as bar plots. A complete list of software packages used for spatial transcriptomic analyses is provided in the Supplementary Table.

### Statistics and reproducibility

Individual data points are displayed in all plots and represent values from independent mice, cells, or biological replicates from cell culture experiments.. Randomization was applied throughout all statistical analyses and in the assignment of mice to treatment groups. Data collection and analysis were not performed in a blinded manner relative to the experiment operators. All analyses, plots, and heatmaps were produced using GraphPad Prism 9.4.1, with statistical significance set at P < 0.05. For t-test and ANOVA analyses, the GraphPad Prism program reports exact P values for P = 0.0001 or higher; for lower values, it reports P < 0.0001.

## Public transcriptomics datasets

Publicly available datasets utilized in this study are listed below: Tsukui et al.(22) (GSE132771), Franzén et al. (29) (S-BSST1410), *Adams et al.(11)* (GSE136831) and *Habermann et al.*(*12*) (GSE1135893).

## Author contributions: Conception and design

K.K., A.A.R.,, X.V., G.L. Analysis and interpretation: K.K., A.A.R., B.S., U.C., A.N., RFN. Drafting the manuscript: K.K., A.A.R., RFN., X.V., G.L. All the authors read and commented on the manuscript

## Acknowledgements

We thank the Boston University single cell RNA-sequencing and flow cytometry core facilities, and support from The Evans Center for Inter-disciplinary Biomedical Research ARC on “Connecting Tissues and Investigators, Fibrosis in Pathology” at Boston University.

## Data Availability

Publicly available datasets utilized in this study are listed below: Tsukui et al.(22) (GSE132771), Franzén et al. (29) (S-BSST1410), *Adams et al.(11)* (GSE136831) and *Habermann et al.*(*12*) (GSE1135893). Newly generated transcriptomic data was deposited to the GEO.

---

**Sup Figure 1.**
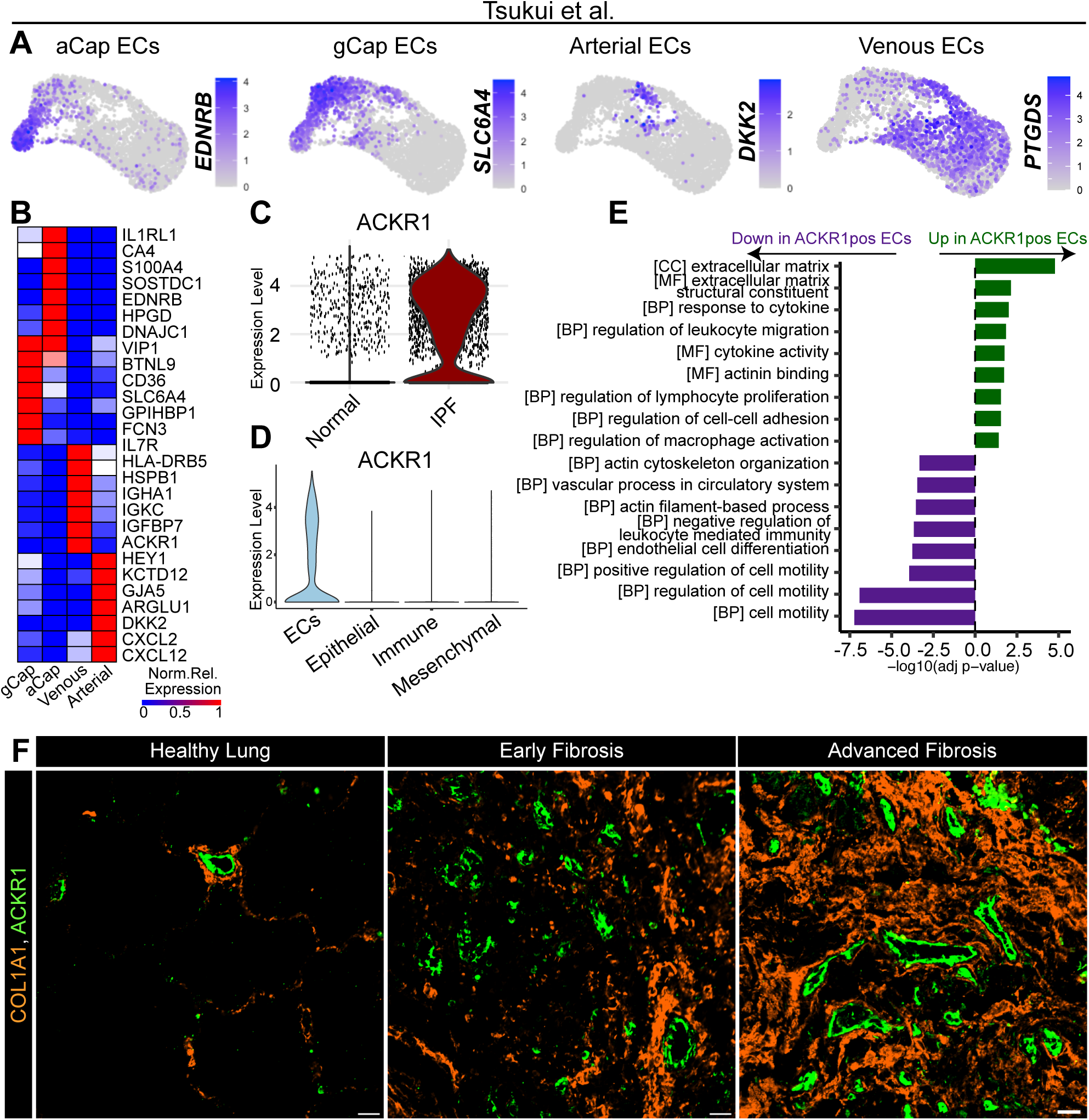
**(A)** UMAP projection of all endothelial cell markers from Tsukui et al. **(B)** Differentially expressed genes between all endothelial subpopulations. **(C)** Violin plot with ACKR1 normalized expression in healthy and IPF endothelial cells. **(D)** Violin plot with ACKR1 normalized expression across all lung lineages. **(E)** GO enrichment of upregulated and downregulated genes.

**Sup Figure 2.**
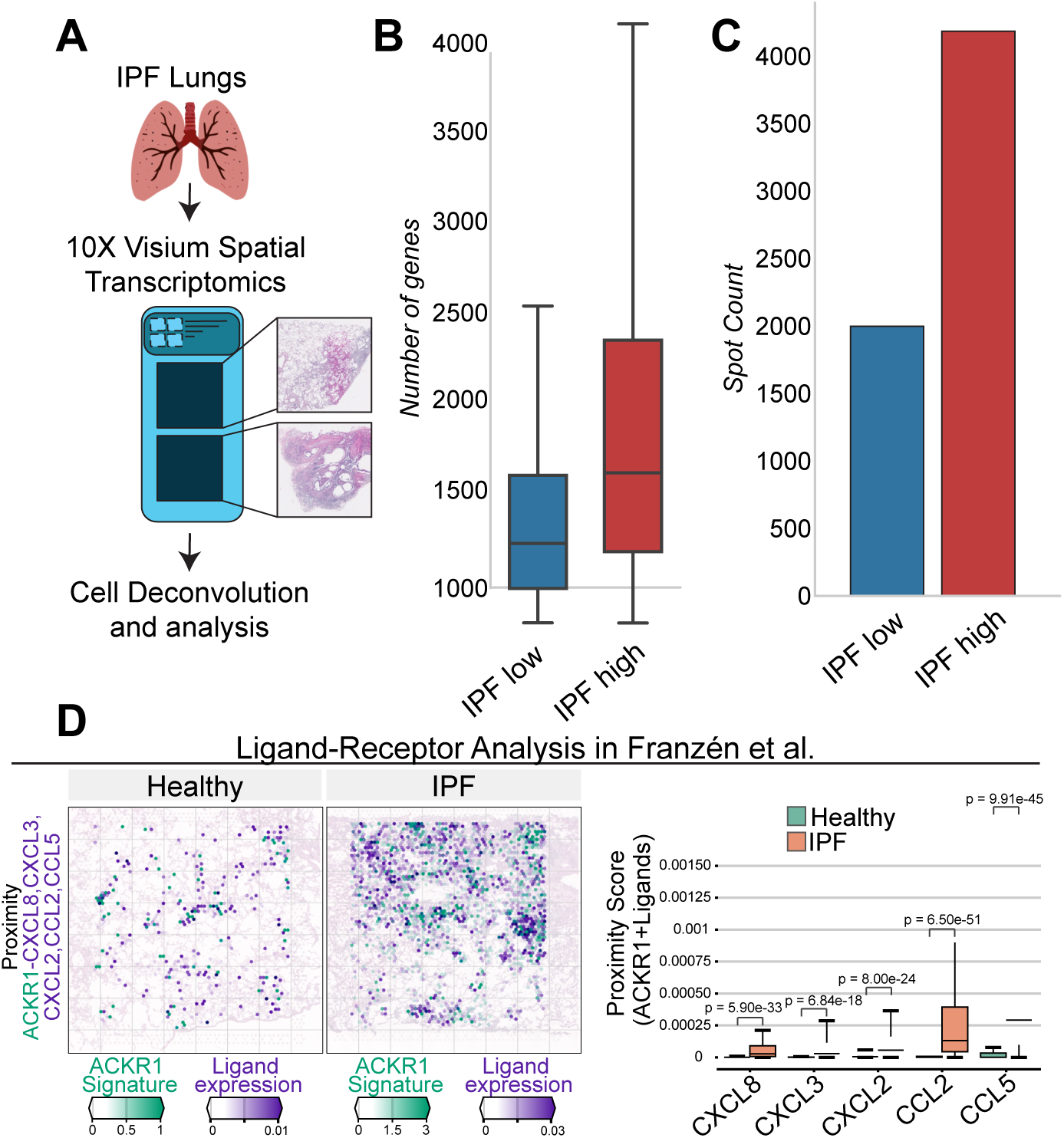
**(A)** Spatial RNA sequencing experimental set up. **(B)** Bar plot of average gene content of spatial RNA sequencing samples. **(C)** Bar plot of spot count of spatial RNA sequencing samples. (**D**) Spatial mapping and quantification of proximity between ACKR1 VECs and ligands in healthy and IPF lungs from the *Franzén et al (29)* dataset. Statistical significance was evaluated using a non-parametric Wilcoxon rank-sum test.

**Sup Figure 3.**
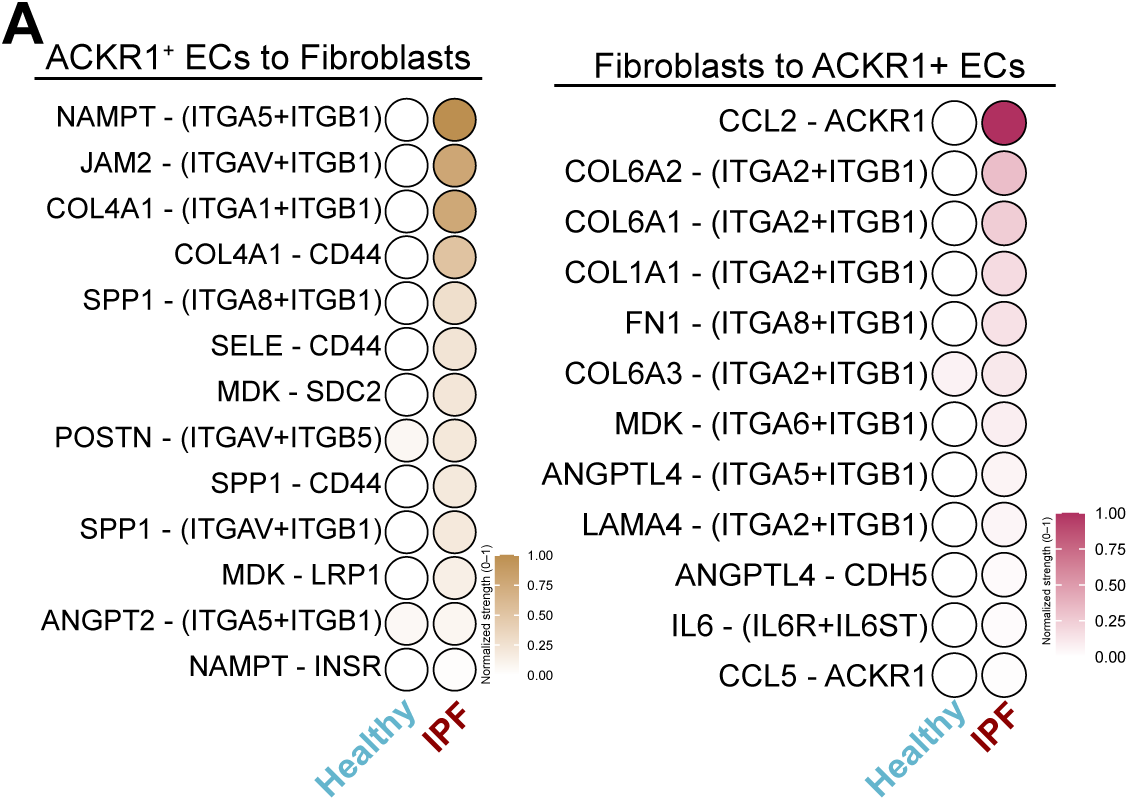
**(A)** Bubble plot with Ligand-Receptor pair interactions and their normalized interaction strength.

**Sup Figure 4.**
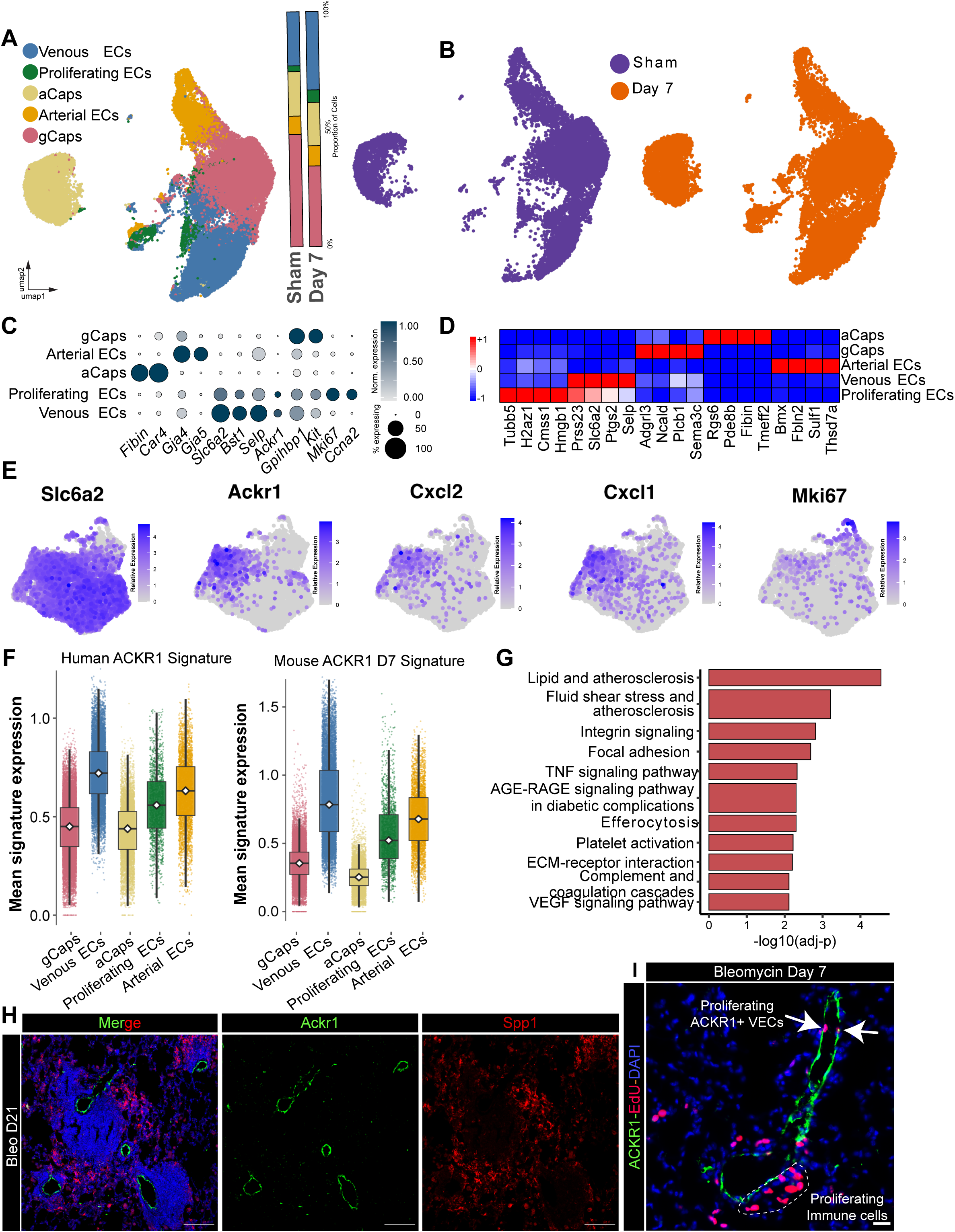
**(A)** UMAP projection of all endothelial cells from sham and day seven post bleomycin and proportion plot. **(B)** UMAP projection of endothelial cells split between sham (purple) and day seven post bleomycin (orange). **(C)** Bubble plot with representative markers for each endothelial lineage. **(D)** Differential gene expression in each endothelial lineage. **(E)** UMAP plots with normalized gene expression across venous endothelial cells **(F)** Human and mouse ACKR1 VEC signature enrichment on all endothelial lineages from sham and bleomycin injured mice. **(G)** KEGG enrichment on mouse ACKR1pos VECs . **(H)** IF for ACKR1 and SPP1 in day day21 mouse bleomycin treated lungs (scale= µm). **(I)** IF for ACKR1 and EdU in day day7 mouse bleomycin treated lungs (scale= 10µm).

